# Expression-System Biases Compromise the Functional Assessment of *KCNH2* Variants and Clinical Risk Assessment

**DOI:** 10.64898/2026.11.04.764433

**Authors:** Aline Rio, Malak Alameh, Sylvain Feliciangeli, Rita Semaan, Marja Steenman, Béatrice Ollivier, Jérôme Montnach, Florian Lesage, Barbara Ribeiro de Oliveira, Michel De Waard, Gildas Loussouarn

## Abstract

**Background:** Functional characterization of *KCNH2* variants is essential for assessing pathogenicity in long QT syndrome type 2. Because hERG channels assemble as tetramers, the relative expression of wild-type (WT) and variant subunits critically influences channel function and pathogenicity assessment. However, commonly used experimental systems may not reproduce the balanced allelic expression observed in heterozygous patients.

**Methods:** Allele-specific *KCNH2* expression was analyzed in two independent RNA-sequencing datasets from human left ventricular tissue. Using the dominant-negative T74R hERG variant as a model, we calibrated the relationship between WT-to-variant expression ratios and hERG current amplitude. Using automated patch-clamp electrophysiology, we compared 3 dual-expression strategies: internal ribosome entry site (IRES), P2A-linked tandem constructs, and dual cytomegalovirus promoter (dCMV) constructs. Co-electroporation of WT and variant plasmids in different ratios was the reference.

**Results:** Allele-specific expression analysis supported balanced *KCNH2* expression in human left ventricular tissue. At a 1:1 WT-to-T74R expression ratio, hERG current amplitude was reduced to 31.0% of WT, consistent with a dominant-negative mechanism in which incorporation of 2 or more T74R subunits renders the channel nonfunctional. All 3 dual-expression strategies produced substantial expression imbalances, with the first cassette consistently favored. IRES constructs showed the greatest asymmetry. P2A constructs partially improved expression balance but introduced an additional N-terminal proline in the second cassette altering T74R-associated trafficking and channel biophysics. The dCMV system also failed to achieve balanced expression, including after stable integration. Co-electroporation of separate WT and variant plasmids provided the most reliable means of controlling their relative expression.

**Conclusions:** Commonly used dual-expression constructs do not reliably reproduce balanced WT and variant hERG expression, potentially distorting assessments of dominant-negative effects and variant pathogenicity. Co-electroporation of separate plasmids provides a robust approach for high-throughput functional phenotyping under heterozygous conditions, with implications for the interpretation of *KCNH2* variants and potentially other genes encoding multimeric ion channels.

## Introduction

Long QT syndrome (LQTS) is an inherited cardiac disorder characterized by prolongation of the QT interval on the electrocardiogram (ECG), which can predispose affected individuals to life-threatening ventricular arrhythmias, including torsades de pointes ^1^. The three major genetic subtypes of LQTS are primarily inherited in an autosomal dominant manner and are associated with variants in *KCNQ1* (LQTS type 1, LQTS1), *KCNH2* (LQTS2), and *SCN5A* (LQTS3). LQTS2 is the second most common subtype accounting for approximately 25-30% of LQTS cases ^2–4^. Inherited LQTS2 is caused by loss-of-function variants in *KCNH2*, which encodes the human ether-à-go-go-related gene (hERG; K_v_11.1) potassium channel responsible for the rapid component of the delayed rectifier potassium current (I_Kr_). In LQTS2, *KCNH2* variants are heterozygous in approximately 99% of cases. The functional hERG channel assembles as tetramers, allowing wild-type (WT) and variant subunits to combine in different stoichiometries. Under the simplifying assumption that only WT and variant subunits are present, five distinct tetrameric compositions are possible, ranging from channels composed entirely of WT subunits to those composed entirely of variant subunits. The relative abundance of these channel populations can substantially influence overall channel function, particularly in the presence of dominant-negative variants that impair the function of WT subunits within the same channel complex ^5^. Consequently, the relative expression levels of WT and variant alleles may critically affect the functional consequences of a given *KCNH2* variant and, therefore, its apparent pathogenicity. Supporting the prevalence of these mechanisms, large-scale multiplexed assay estimated that 42% of hERG variants exhibit at least 50% reduction in protein trafficking, and that 65% of these trafficking-defective variants exert a dominant-negative effect when co-expressed with the WT channel ^6^. Studies using human induced pluripotent stem cell-derived cardiomyocytes (hiPSC-CMs) from patients with LQTS2 have suggested approximately equivalent expression of the WT and variants alleles (50:50) ^7,8^. Although these findings remain to be confirmed in adult cardiac tissues, they provide an important experimental rationale for establishing physiologically relevant WT-to-variant expression ratios when assessing the pathogenicity of *KCNH2* variants.

The widespread adoption of next-generation sequencing has led to an exponential increase in the identification of rare coding variants across the human genome. As of October 2026, ClinVar contains 4019 reported *KCNH2* variants, including 3390 coding variants, of which 1449 are classified as variants of uncertain significance (VUS). Given the potentially fatal consequences of LQTS2, accurate interpretation of hERG variants is essential for patient risk stratification, clinical management, and cascade screening of affected families ^9,10^. According to American College of Medical Genetics and Genomics and the Association for Molecular Pathology (ACMG/AMP) guidelines, well-established functional studies constitutes a source of evidence for variant classification ^11^. With the number of VUS continuing to grow, high-throughput functional phenotyping has become increasingly important for generating the experimental evidence needed to assess the pathogenicity of hERG variants. Several large-scale functional characterization initiatives have been reported to date ^12–16^. However, the reliability of these approaches depends, in part, on whether their experimental models adequately reproduce the heterozygous state, including the relative expression WT and variant alleles. Failure to reproduce this condition may compromise the interpretation of functional data and increase the risk of misclassification. Thus, as the number of VUS characterized by automated patch-clamp (APC) electrophysiology continues to increase, ensuring that experimental models accurately reflect heterozygous expression becomes critical to minimizing the risk of variant misclassification. High-throughput APC studies have adopted different strategies to model the heterozygous condition, including transient expression following electroporation ^17^ and stable expression using bicistronic internal ribosome entry site (IRES)-based constructs ^6,13,15,16,18,19^.

Here, we first investigated allele-specific *KCNH2* expression in adult human left ventricular tissue and demonstrated, for the first time, approximately equal expression of the two alleles. We then sought to identify the experimental strategy that most accurately reproduces this expression balance for the functional assessment of hERG variants under heterozygous conditions. We selected the T74R variant (NM_000238.4.221C>G) as a model because of its pronounced dominant-negative effect on hERG channel trafficking ^5,12^. Using high-throughput APC electrophysiology, we established co-electroporation of separate WT- and T74R-encoding plasmids as a reference approach and compared it with three dual-hERG expression strategies: bicistronic IRES constructs, single-open-reading-frame constructs linking the two hERG sequences through a self-cleaving P2A peptide (P2A), and constructs containing two independent cytomegalovirus (CMV) promoters (dCMV). We assessed the ability of each strategy to reproduce balanced WT and variant expression and to model the functional consequences of heterozygous *KCNH2* variants. Our findings identify co-electroporation as the most reliable of the approaches tested for reproducing balanced WT and variant expression in this experimental system. We confirmed that the pIRES approach underestimates current amplitude under heterozygous condition for 11 additional variants exhibiting a dominant-negative effects. The co-electroporation strategy could therefore provide a robust framework for high-throughput functional characterization of hERG variants and the assessment of their pathogenicity under heterozygous conditions.

## Methods

### Allele-specific expression analysis of *KCNH2*

Allele-specific expression analysis (ASE) of *KCNH2* was analyzed using data from human non-failing left ventricle tissue in two independent RNA-seq datasets: GSE116250 (13 samples) and GSE298265 (6 samples). Analysis was performed following established ASE analysis recommendations ^20^. Reference and alternative allele counts were obtained following WASP filtering to minimize mapping bias ^21^. The alternative allele fraction was calculated as ALT/(REF + ALT). Only SNPs with at least 25 total allele counts per sample were included.

### *KCNH2* plasmid construction

In all the following constructs, we used the reference hERG (K_v_11.1) cDNA (NCBI reference NM_000238.4), modified to add silent restriction sites ^5^.

### Single hERG-encoding plasmids

The pcDNA5/FRT/TO vector expressing WT and T74R was used as described previously ^5^.

### IRES plasmids

Plasmids were generated using a two-step restriction-ligation process on three plasmids: pEX plasmids containing either WT or T74R hERG insert (Eurofins Genomics, Ebersberg, Germany) and the pIRES-EGFP vector (Clontech, Mountain View, CA, USA). The pIRES-EGFP plasmid and the pEX-WT hERG or pEX-T74R hERG insert plasmids were digested at the *XhoI* and *BamHI* sites using high-fidelity enzymes (XhoI-HF® R0146S, BamHI-HF® R3136S, rCutSmart™ Buffer B6004S; New England BioLabs, MA, USA) and inserts were ligated at a 7:1 insert:vector molar ratio (T4 DNA ligase M0202S, New England BioLabs, MA, USA), yielding the intermediate plasmids pIRES-WT-EGFP and pIRES-T74R-EGFP. These intermediates and the pEX-WT hERG or pEX-T74R hERG inserts were subsequently digested at the *BstXI* and *NotI* sites (BstXI-HF® R0113S, NotI-HF® R3189S; New England BioLabs, MA, USA) and ligated to replace GFP by the second hERG (WT or T74R) cassette and obtain the four final plasmid pIRES constructs: WT-IRES-WT, WT-IRES-T74R, T74R-IRES-WT and T74R-IRES-T74R.

### P2A plasmids

WT-P2A-T74R and T74R-P2A-WT plasmids were constructed by synthesizing a P2A sequence followed by a WT or T74R hERG cDNA, containing an introduced *BsrGI* silent site (GenScript Biotech, NJ, USA), and inserting it at the end of the T74R or WT hERG sequence present in a pcDNA5/FRT/TO vector ^5^. To make the two translated proteins as symmetrical as possible and exclude any effect of the C-terminus, the P2A sequence was also added at the end of the hERG of the 2^nd^ cassette, in the initial plasmids sequence. The only remaining difference is that the 2^nd^ cassette hERG starts with a proline, which is unavoidable with the P2A strategy. To produce the other combinations, WT-P2A-WT and T74R-P2A-T74R plasmids, both initial constructions were digested between *BsrGI* and *Xho*I restriction sites using high-fidelity enzymes (BsrGI-HF^®^ R3575S, NotI-HF^®^ R3189S, rCutSmart™ Buffer B6004S, New England BioLabs, MA, USA). WT or T74R inserts were purified on a 0.8% agarose gel and ligated at a 7:1 insert:vector molar ratio (T4 DNA ligase M.202S, New England BioLabs, MA, USA).

### dCMV plasmids

To produce T74R-CMV-WT and WT-CMV-T74R plasmids, WT and T74R hERG cloned in a pcDNA5/FRT/TO plasmid, and two inserts CMV-WT hERG and CMV-T74R hERG were used (Genscript Biotech, NJ, USA). Initial constructs were digested between *MfeI* and *NruI* restriction sites using high-fidelity enzymes (MfeI-HF® R3589S, NruI-HF® R3192S, rCutSmartTM Buffer B6004S, New England BioLabs, MA, USA) and ligated at a 7:1 insert:vector molar ratio (T4 DNA ligase M.202S, New England BioLabs, MA, USA). Both WT-CMV-WT and T74R-CMV-T74R plasmids were then constructed by amplifying overlapping fragments (Q5 Hot start High-Fidelity DNA polymerase M0493S, New England BioLabs, MA, USA) between the restriction sites *HindIII* and *BamHI* (absent in the inserts) using the Gibson assembly strategy (NEBuilder HiFi DNA Assembly Master Mix E2621S, New England BioLabs, MA, USA). For each construct, four primers were designed (**Table 1**). Two external primers overlapped the sequences of two adjacent unique enzyme restriction sites (*HindIII* and *BamHI*) and two internal primers, containing the mutation to be introduced, covering the mutated nucleotide sequence in the forward and reverse directions. Two fragments were thus produced by PCR that overlapped in the region of the inserted mutation and at both extremities to the two adjacent restriction enzymes. Using the relevant plasmid template digested by the selected enzymes and the two mutated amplified fragments, the Gibson assembly process was used to produce the WT-CMV-WT and T74R-CMV-T74R plasmid.

**Table 1.**
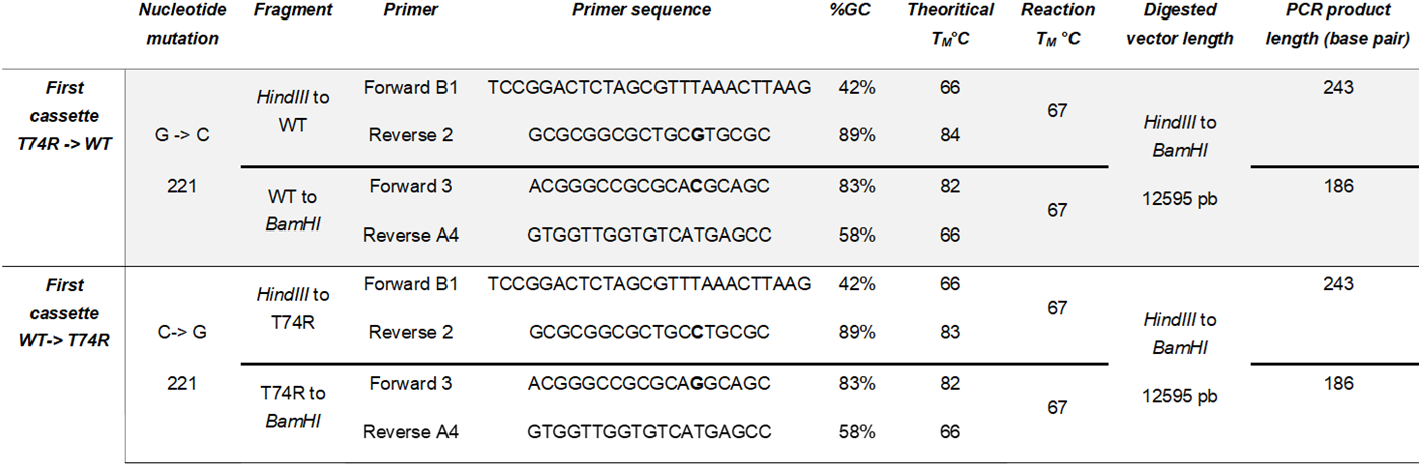
Mutagenesis primers for hERG dCMV plasmids.

All plasmid sequences were confirmed by whole plasmid Nanopore sequencing. Plasmids were purified using the endotoxin-free Qiagen Plasmid Plus Kit (Qiagen, 12945, Hilden, Germany), according to the manufacturer’s protocol, and quantified prior to electroporation or stable cell line preparation.

### Cell culture and transient expression of plasmid

HEK-293T cells (CRL-3216, ATCC) were used for transient expression of the various hERG constructs. Cells were cultured in Dulbecco’s Modified Eagle’s Medium (DMEM; Gibco, France) supplemented with 10% fetal calf serum (Eurobio, PAR, FR), 2 mM L-glutamine, 100 U/mL penicillin, and 100 μg/mL streptomycin (Corning, NY, EUA). Cultures were maintained at 37 °C in a humidified incubator with 5% CO_2_ and regularly tested for mycoplasma contamination using MycoAlert (Lonza, Rockland, ME, USA). Each plasmid was introduced into HEK-293T cells via electroporation using OC-100 cassettes with the MaxCyte STx system (MaxCyte Inc., MD, USA), as previously described ^5^. In conditions in which a single plasmid was transfected (IRES, P2A, dCMV), 24 μg of DNA were used, together with 1 μg of eGFP plasmid, for a total of 25 μg of DNA. In conditions in which two hERG plasmids (WT and T74R) were transfected, different proportions of WT hERG DNA (pcDNA5 WT) and variant hERG DNA (pcDNA5 T74R) were co-transfected, also with 1 µg of eGFP, to achieve a total DNA concentration of 25 μg (**Table 2**).

**Table 2.**
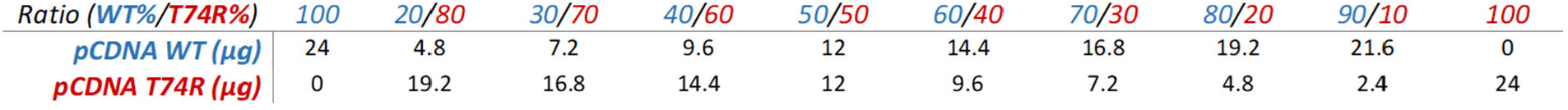
Different proportions of hERG WT DNA (pcDNA WT hERG) and hERG variant DNA (pcDNA5 T74R hERG).

Thirty hours post-electroporation, cells were detached using Accutase (Sigma-Aldrich, St. Louis, MO, USA) and resuspended for APC assays.

### Generation and culture of stable inducible cell lines

Stable, tetracycline-inducible cell lines were generated using the Flp-In T-REx system (Invitrogen, MA, USA) according to the manufacturer’s recommendations. HEK293 Flp-In T-REx host cells were maintained under standard culture conditions in complete growth medium (as described above), supplemented with blasticidin (15 µg/mL) and zeocin (100 µg/mL). Cells were co-transfected at a 1:9 ratio with the pcDNA5/FRT/TO expression plasmid and the pOG44 plasmid encoding the Flp recombinase using Lipofectamine 2000 (Invitrogen, MA, USA) following the manufacturer’s instructions. Forty-eight hrs post-transfection, cells were split and replated at 25% confluence, and zeocin was replaced with hygromycin B (150 µg/mL). Selection medium was refreshed regularly until resistant cell populations were established. Hygromycin-resistant clones were pooled. Expression of the recombinant protein was confirmed by electrophysiological recording. A dose-response was performed to determine tetracycline concentration for optimal induction.

### High-throughput APC recordings

Evaluation of the different heterozygous expression conditions was performed using an optimized fast-track protocol on the SyncroPatch 384PE system (Nanion, Munich, Germany), as previously described ^22^. Recordings were performed using an external NMDG-based solution (in mM: 80 NaCl, 4 KCl, 2 CaCl_2_, 1 MgCl_2_, 5 glucose, 60 N-Methyl-D-Glucamine (NMDG), 10 HEPES; pH 7.4 adjusted with NaOH; 280 ± 3 mOsm) and an internal solution (in mM: 110 KF, 10 NaCl, 10 KCl, 10 EGTA, 10 HEPES (pH 7.2, NaOH), 280 ± 3 mOsm). Briefly, the holding potential was set at −86 mV, and step durations were adjusted to reach steady-state based on the voltage-dependent kinetics of WT and T74R channels. The final fast-track protocol consists of six sub-protocols that allow the recording, in the following order, of steady-state activation, steady-state inactivation, deactivation, inactivation, recovery from inactivation, and activation kinetics. The complete protocol lasts 74 sec. Maximal currents were measured using the following voltage protocol: a 2-s depolarization step to +60mV to open the activation gate, a 10-ms repolarization step to −120mV to open the inactivation gate, and a test step at +60mV. We tested various heterozygous ratios of single hERG-encoding plasmids (WT *versus* T74R) via co-electroporation to calibrate the degree of negative dominance of the T74R variant. We then compared the various heterozygous expression strategies, P2A, IRES, dCMV, each with four order combinations (WT-WT; WT-T74R; T74R-WT; T74R-T74R). Each construct was tested in two replicates.

### Modeling the dominant-negative effect of a variant on the hERG channel

To model the dominant-negative effect of a mutant subunit in a tetrameric potassium channel, we used a binomial distribution assuming random assembly of WT and variant subunits. The probability of forming a tetramer containing *k* variant subunits was calculated as:

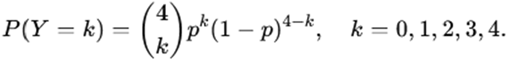

where (p=X/100), and *X* represents the percentage of variant subunits expressed (ranging from 0 to 100%).

### Automated data analysis and statistical analyses

All R scripts used to analyze the APC data are available at https://github.com/jeromemontnach/hERGvariants. The following quality controls were used to exclude cells for data analyses: access resistance Ra > 10 MΩ, seal resistance Rs < 600 MΩ measured at −86 mV, leak currents amplitude >200 pA at −60 mV, or run-down >20% over the course of the experiment. For the extraction of biophysical properties, a minimal current amplitude of 500 pA was required, in order to address hERG channel populations with properties unaffected by the contribution of native K^+^ channels in HEK-293T cells (when <500 pA), we also excluded cells with amplitudes above 10 nA to avoid recordings with poor voltage control ^23^. Statistical analyses were performed using non-parametric Mann-Whitney t-tests for current amplitudes and two-way ANOVA. Significance thresholds were set as follows: p < 0.05 (*), p < 0.01 (**), p < 0.001 (***), p < 0.0001 (****).

## Results

### Allelic expression of *KCNH2* in adult human ventricular tissue

To quantify allelic expression of *KCNH2* in human left ventricular tissue, alternative allele (ALT) expression ratios were assessed using several heterozygous SNPs distributed across the gene in 13 and 6 human left ventricular samples from two independent RNA-seq datasets. For each SNP, the ALT allele fraction was calculated as the proportion of transcripts derived from the alternative allele relative to the total number of transcripts derived from both alleles. A theoretical ratio of 0.5 indicates balanced allelic expression. Across the five SNP positions analyzed, median ALT allele fractions ranged from 0.36 (SNP 150951701) to 0.70 (SNP 150951110) in the first dataset (GSE116250; **Figure 1**) and from 0.43 (SNP 150948446) to 0.59 (SNP 150952515) in the second dataset (GSE298265; **Figure S1**). Overall, the observed values, which were generally close to 0.5, support broadly balanced *KCNH2* allelic expression in human left ventricular tissue across both RNA-seq datasets.

**Figure 1:**
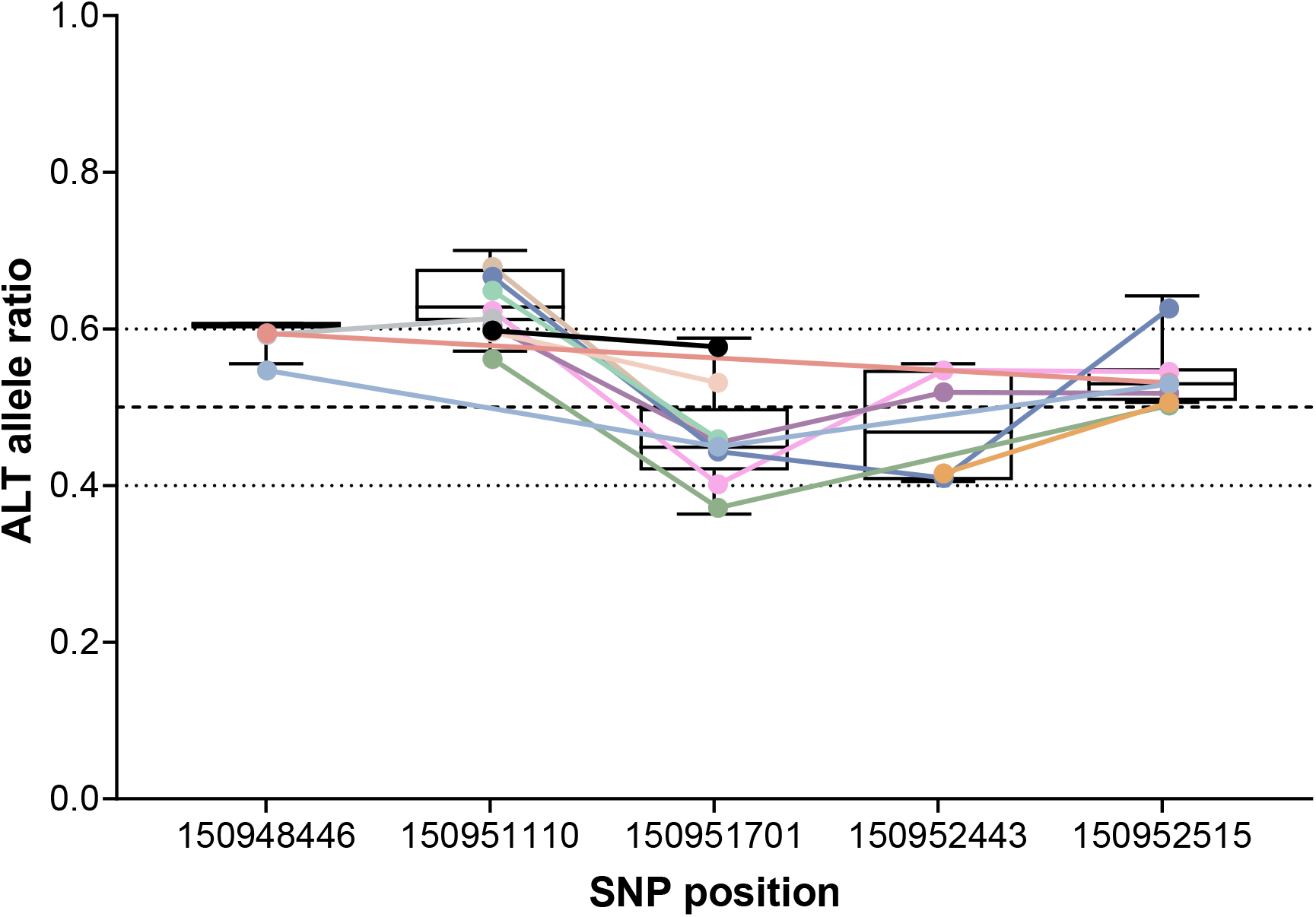
Allele-specific expression of *KCNH2* in the GSE11625 RNA-seq dataset. Alternative allele fractions (ALT/[REF + ALT]) are shown for individual SNPs across samples. Only SNP-sample observations with at least 25 total allele counts were included. Colored points represent individual samples, with lines connecting SNP measurements from the same sample. Boxplots show the distribution of allelic fractions at each SNP. The dashed horizontal line indicates balanced allelic expression (0.5), while dotted lines indicate reference values of 0.4 and 0.6. ^24^

### Theoretical considerations on the dominant-negative effects of hERG variants

The severity of a dominant-negative effect depends on the minimum number of WT subunits required for a tetrameric channel to remain functional. Four main models of dominant-negative effects can be distinguished, depending on the number of variant subunits that can be accommodated within a functional tetramer, with each model predicting a different degree of functional impairment (**Figure 2A**). As the relative expression of the variant increases compared with that of WT hERG, the equilibrium shifts towards the formation of tetramers containing a greater number of variant subunits, ultimately favoring tetramers composed entirely of variant subunits. To model this phenomenon, we used a binomial distribution of variant assembly to generate four theoretical curves describing the normalized current amplitude as a function of the proportion of expressed variant subunits (**Figure 2B**).

**Figure 2:**
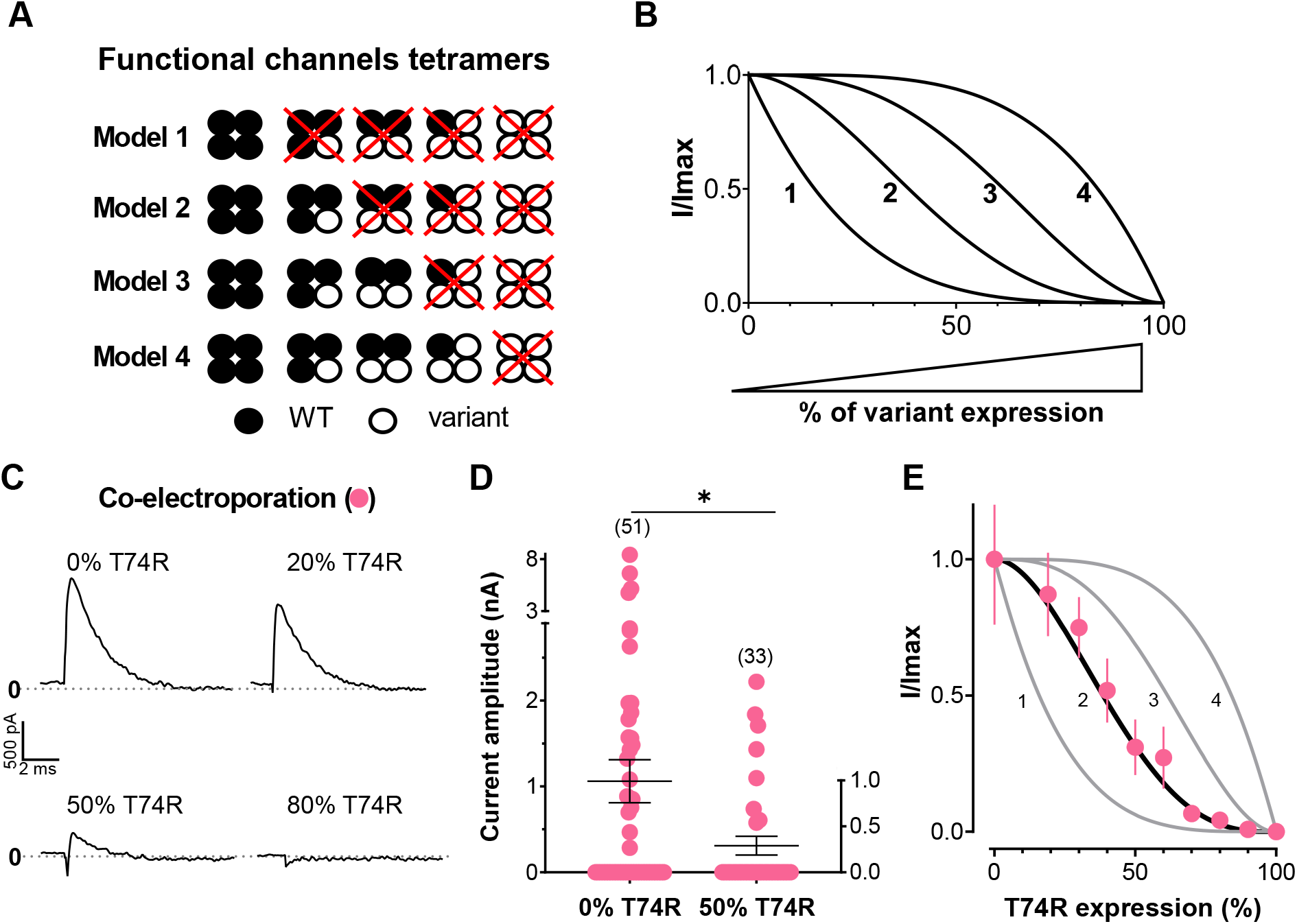
Theorical and experimental dominant negative effects of variants on WT/T74R hERG channel function, modeled with a binomial model. **A)** All possible combinations of WT and mutant subunits within the tetramer for each model of negative dominance, with WT subunits in black and variant in white. Red crosses indicate non-functional channels in the various models 1 to 4. **B)** Theoretical curves representing, in the different models, the relative hERG current as a function of the proportion of mutant subunits (%). **C)** Representative current recordings in HEK-293 cells co-transfected with 4 different ratio of WT hERG and T74R variant. **D)** Current amplitudes observed in each cell for different ratios of co-transfected WT and T74R plasmids. Data are presented as mean ± SEM. *p<0.05 test, Mann-Whitney t-tests. **E)** Normalized experimental data obtained from WT:T74R co-transfection ratios, overlaid on the binomial dominant-negative model, resulting in a theoretical calibration curve. Data are presented as mean ± SEM.

### Experimental calibration of the dominant negative effect of the T74R variant

To determine which model best describes the dominant-negative effect of the T74R variant, we co-expressed WT and T74R hERG using the corresponding single-gene expression plasmids at different relative proportions. The proportion of variant-encoding plasmid was varied from 0 to 100% (**Table 2**), and hERG currents were measured by APC. Representative current traces illustrate the reduction in current amplitude as the proportion of T74R-encoding plasmid increases (**Figure 2C**). The resulting mean normalized current amplitudes were then compared with the four theoretical curves representing the different models of negative dominance (**Figure 2E**). The experimental data closely matched Model 2 (**Figure 2A**), in which a tetramer can retain normal channel activity in the presence of a single variant subunit, whereas the incorporation of two or more variant subunits abolishes channel activity. Under the 50% T74R / 50% WT condition, corresponding to an equal WT-to-variant expression ratio, the current amplitude decreased to 31.0% of the WT current, closely matching the theoretical value of 31.3% predicted by Model 2 (**Figure 2D)**. This result is consistent with the previously reported dominant-negative effect of T74R ^5,12^. Under this perfectly balanced expression ratio, T74R changed neither activation nor inactivation at steady-state, only kinetics were slightly altered, compared with WT channels (**Figure S2B**). This suggests that T74R effect is primarily attributable to impaired channel trafficking^5^. Having established the dominant-negative model that best describes T74R and derived an experimental calibration curve, we then used this framework to assess the relative expression of WT and T74R subunits in the IRES-, P2A-, and dCMV-based expression systems.

### Estimation of WT-to-T74R hERG expression ratios using the IRES system

We next evaluated bicistronic constructs incorporating an IRES to assess the relative expression of WT and T74R hERG channels. The IRES-based dual-hERG expression system enables the expression of two hERG channels from a single mRNA, with the two coding sequences separated by an IRES. Translation of the first coding sequence is cap-dependent: eIF4F recognizes the 5′ cap, recruits the 40S ribosomal initiation complex, and enables ribosomal scanning to an AUG start codon within a favorable initiation context ^25,26^. In contrast, translation of the downstream coding sequence depends on the IRES-mediated internal recruitment of ribosomes, a generally less efficient process ^27^. To estimate the relative expression of WT and T74R channels, we generated four IRES constructs (IRES WT-WT, IRES T74R-T74R, IRES WT-T74R, and IRES T74R-WT). These constructs allowed us to assess current amplitudes according to channel composition (WT or T74R) and the relative position of the WT and variant coding sequences within the bicistronic construct. Representative current traces showed the expected maximal current amplitude for IRES WT-WT and the absence of detectable current for IRES T74R-T74R (**Figure 3A**). Notably, the IRES T74R-WT construct exhibited a complete loss of detectable current despite the presence of the WT coding sequence, consistent with a strong dominant-negative effect in this order. Conversely, the IRES WT-T74R construct produced currents close to those recorded with IRES WT-WT in spite of the presence of the T74R in the second cassette (**Figure 3A**). Population-level recordings confirmed this trend, with only a modest, statistically nonsignificant reduction in current amplitude for IRES WT-T74R (**Figure 3B**). In contrast, no detectable current was recorded with the IRES T74R-WT construct. Thus, the two constructs, which differed only in the relative order of the WT and T74R coding sequences, exhibited markedly different functional outcomes (p < 0.0001), confirming that cassette position substantially influences the relative expression of the two channels. To estimate the relative expression of T74R, we plotted the percentage of residual current against the theoretical model 2 of dominant-negative effects. With the IRES WT-T74R construct, the mean residual current was 78%, whereas no residual current was detectable with IRES T74R-WT (**Figure 3C**). According to the theoretical model, these results correspond to a maximum T74R expression fraction of 22.5% for IRES WT-T74R, implying at least a 3.5-fold higher expression of WT than T74R. Conversely, the absence of detectable current with IRES T74R-WT suggests that T74R accounts for more than 90% of the expressed channels, corresponding to a greater than 9-fold excess of T74R over WT. These findings raise fundamental concerns about the use of IRES-based dual-hERG expression systems in which the variant coding sequence precedes the WT sequence. Under these conditions, the resulting expression imbalance may approach a functionally homozygous state, in which the variant channel overwhelmingly predominates despite the presence of a WT coding sequence. This limitation is particularly important when such systems are used to investigate dominant-negative effects or model the heterozygous state associated with autosomal dominant *KCNH2*-related disease. Finally, consistent with the data generated by the co-electroporation of two independent plasmids, T74R expression (through the second cassette of the IRES-based dual-hERG expression system) did not measurably alter the biophysical properties (**Figure S3**).

**Figure 3:**
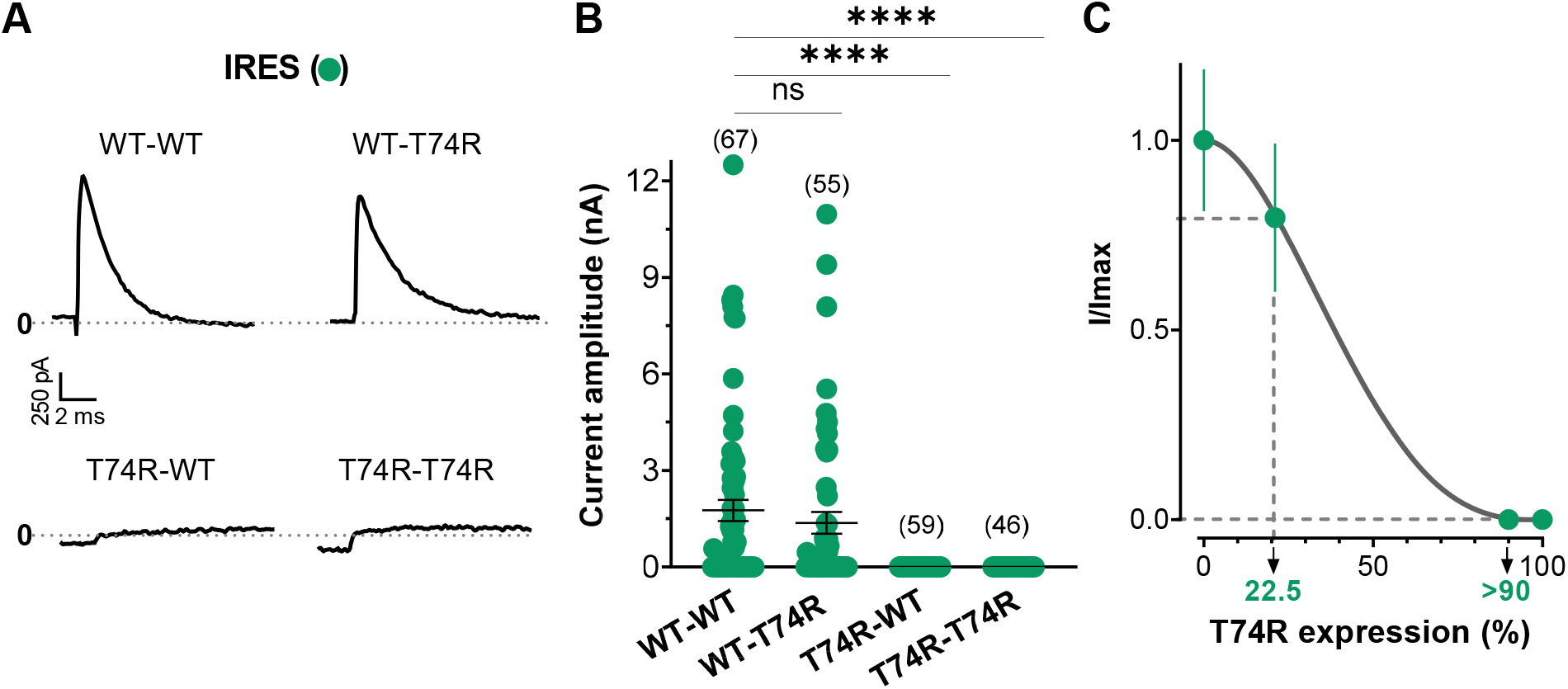
Construct order shapes the relative WT/mutant expression ratio in IRES bicistronic systems. **A)** Representative current recordings in HEK-293 cells transfected with the 4 pIRES plasmids. **B)** Current amplitudes of different pIRES plasmids. Data are presented as mean ± SEM. ****p<0.0001. **C)** Normalized current amplitude data obtained from pIRES plasmid transiently expressed, overlaid on the binomial dominant-negative model #2.

### Efforts to optimize dual-hERG expression systems for balanced WT and variant expression

Although our results indicated that the IRES bicistronic expression system was unsuitable for achieving equivalent expression of WT and variant hERG channels, we investigated alternative dual-hERG expression strategies. We first revisited the P2A expression system using four constructs: P2A WT-pWT, P2A T74R-pT74R, P2A WT-pT74R, and P2A T74R-pWT (p standing for proline). In this bicistronic design, the two hERG expression cassettes are separated by a porcine teschovirus-1 2A (P2A) sequence comprising 21 amino acids and ending in a Gly-Pro motif. A single mRNA is translated continuously from a single initiation site, with the P2A sequence inducing ribosomal skipping between the terminal glycine and proline residues. This mechanism allows the production of two separate proteins, theoretically at near-equimolar levels. To minimize differences between the two translated proteins, the 21 amino acids preceding the terminal Gly-Pro motif were also added to the C-terminus of the hERG protein encoded by the second cassette, ensuring that both proteins carried the same C-terminal extension (**Figure 4A**). The only remaining difference was the presence of an additional N-terminal proline residue immediately preceding the methionine of the hERG protein encoded by the second cassette, an unavoidable consequence of the P2A strategy. Representative current traces revealed a pattern similar to that observed with the IRES expression system (**Figure 4B**). The P2A T74R-pWT construct exhibited markedly reduced current amplitudes compared with P2A WT-pWT, consistent with a strong dominant-negative effect. However, unlike the IRES T74R-WT construct, residual currents remained detectable, suggesting that WT channels were better expressed in the P2A system. Surprisingly, the P2A WT-pT74R construct did not exhibit any reduction in current amplitude, indicating that the expected dominant-negative effect of the T74R variant was no longer apparent. These observations were confirmed by population-level recordings, which showed an 82% reduction in mean current amplitude for the P2A T74R-pWT construct relative to P2A WT-pWT (**Figure 4C**). We hypothesized that the loss of the dominant-negative effect observed with P2A WT-pT74R might result from the additional proline residue at the N-terminus of the T74R-containing hERG protein, which is absent from the P2A T74R-pWT construct. This additional residue might partially compensate for the trafficking defect associated with T74R. We then mapped the residual current observed with P2A T74R-pWT (18% of the current measured with P2A WT-pWT) onto theoretical model 2 of dominant-negative effects (**Figure 4D**). According to this model, the observed current reduction would correspond to a T74R subunit fraction of approximately 60%, with WT subunits accounting for the remaining 40%. Thus, the model predicts a 1.5-fold higher abundance of variant than WT subunits in the P2A T74R-pWT configuration. This analysis could not be applied to the P2A WT-pT74R construct because no current inhibition was observed relative to P2A WT-pWT. Consistent with a potential effect of the additional N-terminal proline residue, we also detected a change in deactivation, slower in hERG channels expressed using P2A WT-pT74R (despite low expression of T74R) that was absent from the reciprocal P2A T74R-pWT construct, despite high expression of T74R (**Figure S4C**). Also surprisingly, half-inactivation potential was shifted toward more positive values in hERG channels expressed using the WT-pWT (compare with **Figures S2C, S3C, S5C, S6D**) and T74R-pWT constructs, but not the reciprocal WT-pT74R construct, suggesting that the introduced N-terminal proline residue affects only the WT but not the T74R subunit conformation linked to inactivation. Taken together, these findings suggest that the P2A dual-hERG expression system could represent an interesting alternative to the IRES system for achieving balanced expression of WT and variant channels, provided that the variant is placed in the first expression cassette, as is commonly done with IRES-based constructs ^28,29^. However, our results also highlight the potential pitfalls of introducing non-native amino acids into hERG proteins when assessing the pathogenic effects of variants. In this case, the addition of a single proline residue was associated with the apparent loss of the dominant-negative effect of T74R and major biophysical alterations.

**Figure 4:**
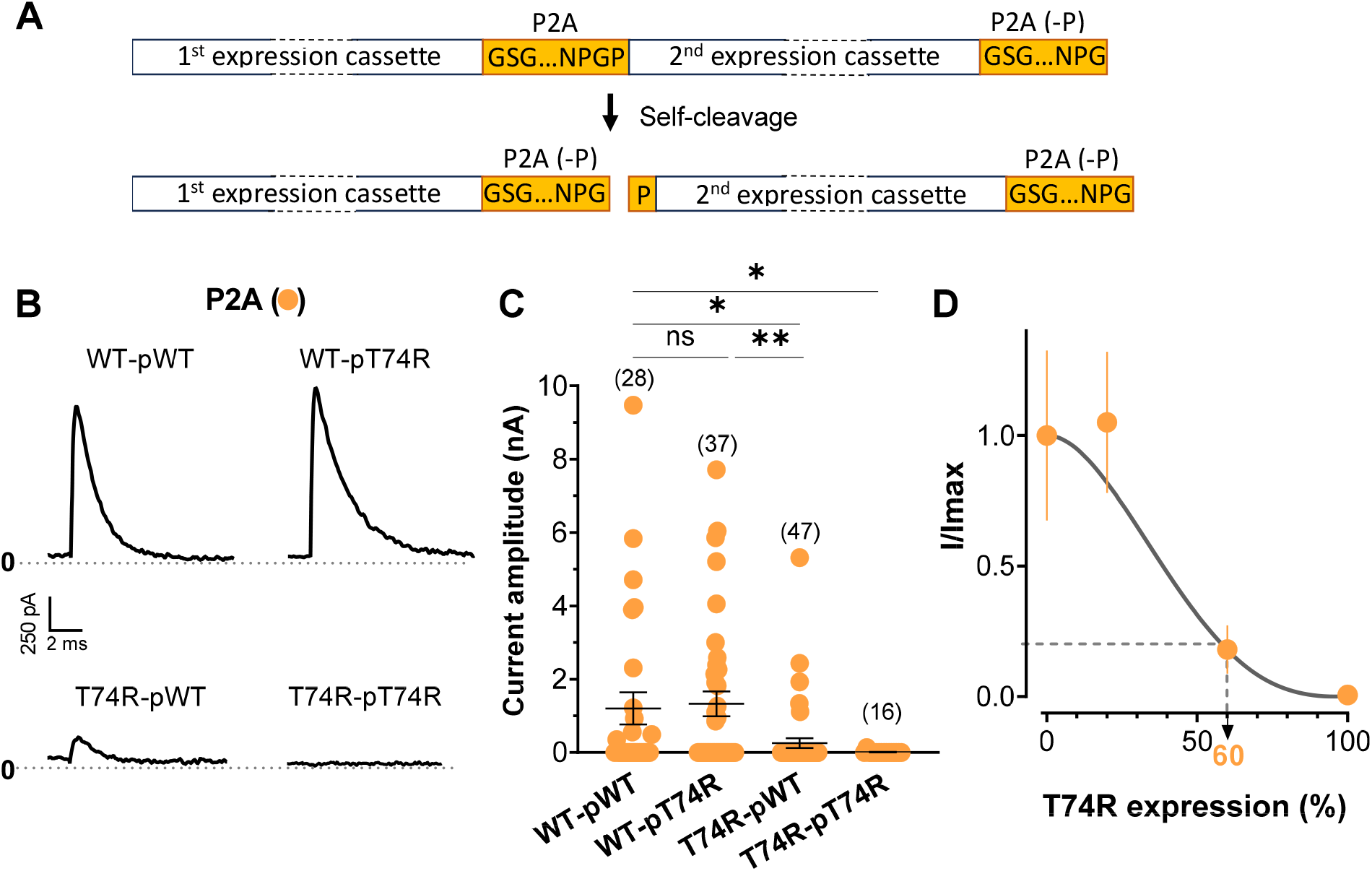
Construct order shapes the relative WT/mutant expression ratio in P2A bicistronic systems. **A)** Schematic representation of cleavage occurring in P2A plasmid construction. The arrowhead indicates self-cleavage site. **B)** Representative current recordings in HEK-293 cells transfected with 4 P2A plasmids. B) Current amplitudes of different P2A plasmids. Data are presented as mean ± SEM. *p<0.05, **p<0.01 **C)** Normalized current amplitude data obtained from P2A plasmid transiently expressed, overlaid on the binomial dominant-negative model #2.

As another strategy to achieve balanced WT and variant expression, we designed a plasmid containing two independent expression cassettes, each driven by its own CMV promoter. Because both cassettes were under the control of identical promoters, we hypothesized that this dual-promoter design would support equivalent expression of WT and T74R channels. An additional advantage over the P2A system was that it did not introduce any non-native amino acids into the hERG sequence, as was also the case with the IRES system. Representative current traces revealed a pattern broadly similar to those observed with the IRES and P2A expression systems (**Figure 5A**). Importantly, both dCMV WT-T74R and dCMV T74R-WT exhibited dominant-negative effects of T74R. Surprisingly, however, cassette order continued to influence the magnitude of current suppression, with the variant in the first cassette producing the strongest effect. Population-level recordings confirmed these observations, showing a mean current reduction of 41% for dCMV WT-T74R relative to dCMV WT-WT, compared with a 91% reduction for dCMV T74R-WT (**Figure 5B**). We next mapped the residual currents measured with the dCMV WT-T74R and dCMV T74R-WT constructs, relative to dCMV WT-WT, onto theoretical model 2 of dominant-negative effects (**Figure 5C**). According to this model, the T74R subunit fraction was estimated at approximately 32% for dCMV WT-T74R and 68% for dCMV T74R-WT. As expected, the 32% T74R expression level did not affect the biophysical properties of the expressed hERG channels (**Figure S5**). These results indicate that the dCMV dual-hERG expression system performed equivalently when the variant was placed in the first cassette (T74R-WT) or in the second cassette (WT-T74R): there was the same 18% difference with the targeted 50%, the first construct over-expressing, and the second construct under-expressing the variant. Nevertheless, the estimated T74R subunit fraction in the dCMV T74R-WT configuration (68%) was slightly higher than that predicted for the P2A T74R-pWT construct (60%). Thus, despite the use of two independent promoters, the dCMV system did not provide any improvement over the P2A approach. It remains possible that additional factors influenced their relative expression levels. In particular, the large plasmid size (13,024 bp) and the presence of two similar CMV-hERG cassettes within the same supercoiled circular DNA molecule may have affected the relative accessibility of the two transcriptional units, as suggested earlier. Previous work has shown that plasmid topology and the structural organization of supercoiled plasmids can influence transgene expression following transfection ^30^. One possibility is that the first cassette was more transcriptionally accessible than the second, resulting in higher expression from the former. To investigate whether plasmid topology and configuration contributed to this imbalance, we generated stable HEK293 cell lines by integrating a linearized version of the construct using the Flp-In/FRT system. However, current amplitudes remained strongly dependent on cassette orientation in these stable cell lines (**Figure S6A**). Indeed, the imbalance was even more pronounced: the dCMV WT-T74R construct no longer produced detectable current suppression, whereas the dCMV T74R-WT construct produced a strong dominant-negative effect, with a mean current reduction of 93.8%, comparable to the 91% reduction observed under transient-expression conditions. Thus, stable integration completely abolished the detectable dominant-negative effect of T74R when the variant was encoded by the second cassette, suggesting that its expression was insufficient to exert a measurable functional effect in this configuration. Overall, stable integration did not improve the balance between WT and variant expression. These findings further demonstrate that neither the use of two independent CMV promoters nor stable integration of the expression construct was sufficient to ensure equivalent expression of WT and variant hERG channels.

**Figure 5:**
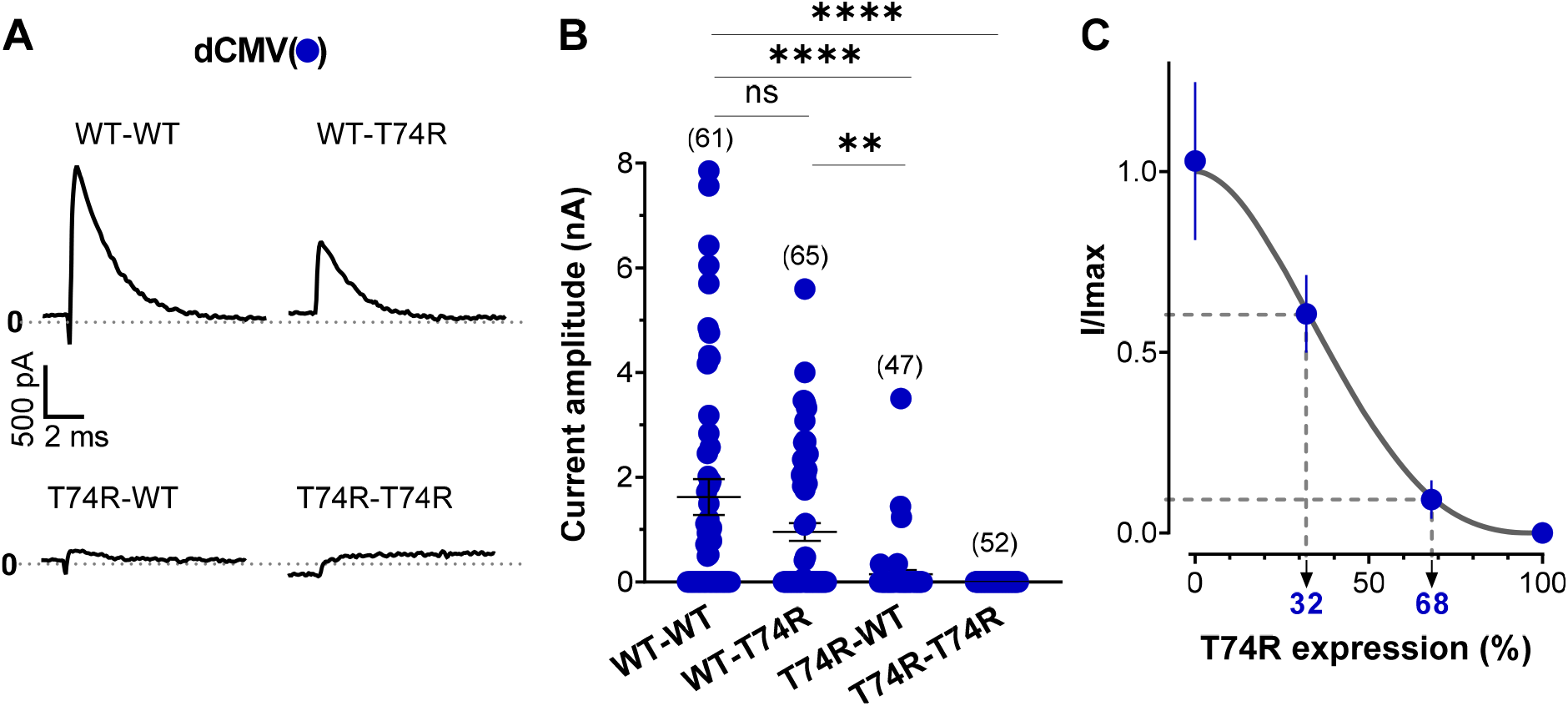
hERG current amplitude from transient expression of dual-CMV systems reveals the relative WT/mutant expression. **A)** Representative current recordings in HEK-293 cells transfected with 4 dual-CMV plasmids. **B)** Current amplitudes of different dual CMV plasmids transiently expressed. Data are presented as mean ± SEM. **p<0.01; ****p<0.0001. **C)** Normalized current amplitude data obtained from transiently expressed dual CMV plasmids, overlaid on the binomial dominant-negative model #2. The experimental data suggest an approximate expression ratio of 68%/32% for the dual CMV system.

### The negative consequence of using the IRES system can be generalized to several other hERG variants

Given that the use of the IRES strategy has been generalized in several high-throughput phenotyping studies ^6,13,15,16,18,19^, we further investigated whether the unbalanced expression observed with the IRES system for the T74R variant is also true for other hERG dominant negative variants. For that purpose, we compared data generated by APC in heterozygous condition using either the co-electroporation method^31^ or the IRES system^18^. Since other variants should not necessarily obey to Model 2 of negative-dominance, we plotted theoretical current amplitude across the entire spectrum of negative dominance (**Figure 6A**,**B**). Model 1 being the strongest negative-dominance, and model 4 being the weakest, they differ in the percentage of remaining current. If both expression systems were equivalent, we should observe the experimental data onto 1X blue line (**Figure 6B**). On the contrary, what was observed, is that the IRES system amplified the negative dominance of most variants. Fitting the data indicate an average of 3.5x overexpression of the variant over the WT with the IRES system (**Figure 6B**). For comparison, a 9X curve is also illustrated. Altogether, the average overestimated decrease in current amplitude by IRES negative dominance reaches 66.4 +/-5.9% (**Figure 6C**).

**Figure 6:**
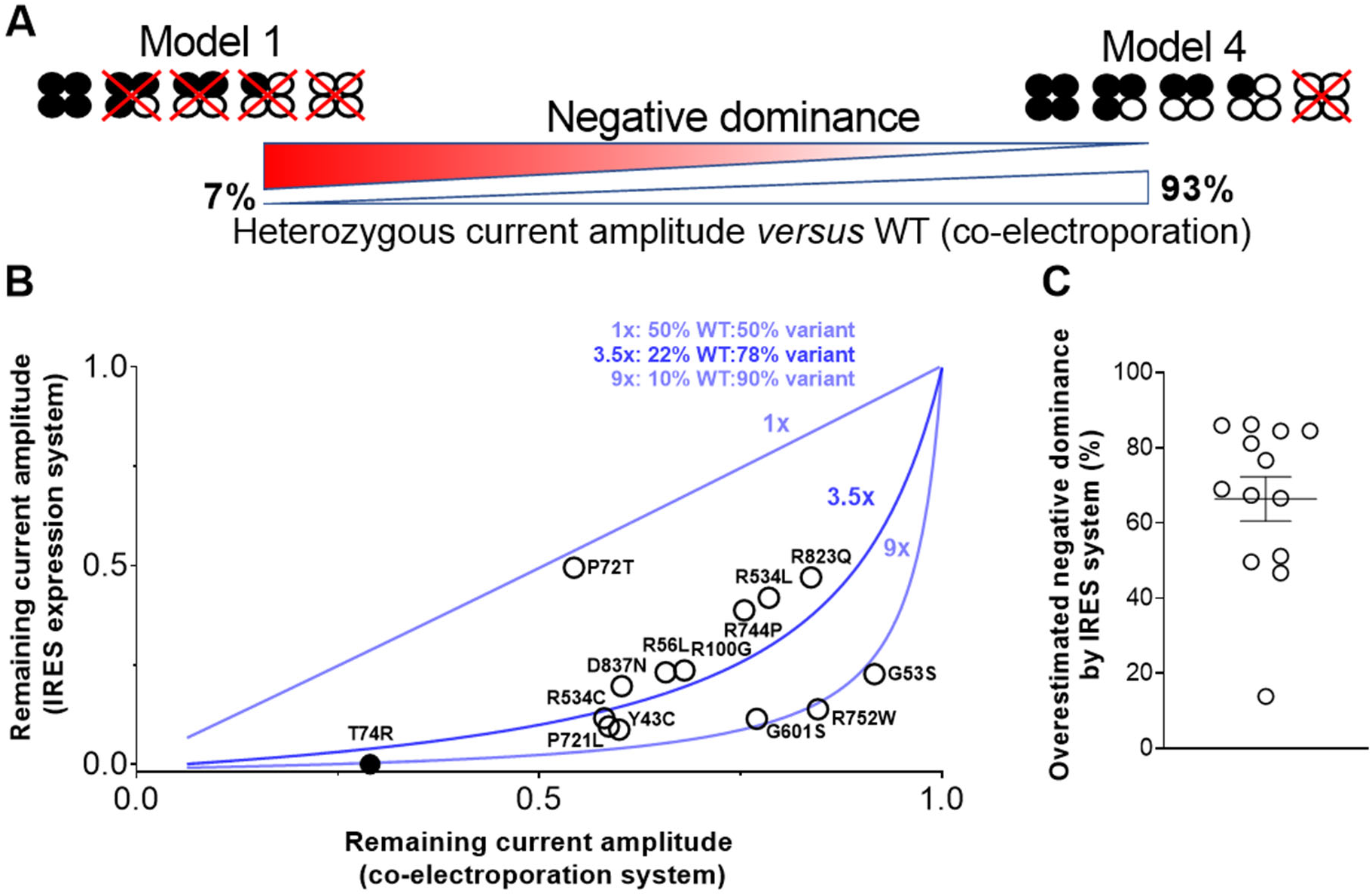
Heterozygous hERG current amplitude relative to WT: Comparison of LQT2 variants studied in two cellular models of heterozygous expression: bicistronic pIRES versus co-electroporation of pcDNA5. **A)** Variants selected here are all trafficking defects variants both studied in Ribeiro de Oliveira et al, 2026 ^31^ and O’Neill et al, 2024 ^18^. Decreasing negative dominance (model 1 to model 4) leads to higher relative currents. Percentages corresponds to min and max current amplitude due to binomial distribution. **B)** Most of the variants (13/14) give rise to lower relative current in pIRES model relative to pcDNA5 co-electroporation model, consistent with an overexpression of the variant (first cassette) as compared to WT (second cassette). Lines: mathematic models of equal expression (1X), 3.5 and 9-fold expression of the first cassette as compared to the second cassette. **C)** Extent of negative dominance bias induced by the IRES system on all variants common to Ribeiro de Oliveira et al, 2026 and O’Neill et al, 2024.

## Discussion

With the launch of several high-throughput automated patch-clamp phenotyping projects to assess the pathogenicity of hundreds of ion channel variants, the use of reliable expression systems has become essential. The aim of this study was to compare different expression systems in terms of their ability to reproduce equal allelic expression. This is particularly important for autosomal dominant diseases such as LQTS2, which is caused by pathogenic *KCNH2* variants. Failure to maintain an appropriate balance between WT and variant channel expression may lead to erroneous conclusions, particularly when assessing variants of VUS with dominant-negative effects. Here, we demonstrate that equal allelic expression of *KCNH2* is the rule in the human heart, highlighting the need for experimental expression systems to reproduce this physiological condition. Our systematic comparison yielded several important conclusions.

First, the IRES system, which is widely used in the literature, proved to be the least suitable approach for achieving balanced expression of WT and variant hERG channels under heterozygous conditions. Placing the variant in the first cassette resulted in its expression reaching levels up to ninefold higher than those of the WT channel. This substantial imbalance is likely to overestimate the dominant-negative effect and makes the experimental condition more closely resemble the homozygous state than the heterozygous condition encountered in patients. These findings are consistent with previous observations showing that expression from the second cassette is approximately 3- to 7-fold lower than that from the first cassette ^28^.

Second, the first cassette consistently exhibited higher expression than the second across all dual-hERG expression systems tested. Among these systems, the P2A variant-WT construct provided the closest approximation to equal allelic expression, with only a 1.5-fold higher expression level of the variant channel. Although this imbalance is substantially smaller than that observed with the IRES system, it remains a deviation from the physiological 1:1 expression ratio.

Third, both the IRES and dCMV systems preserve the integrity of the hERG amino acid sequence, as expected. In contrast, the P2A system introduces additional amino acids at both channel termini: an extra proline residue at the N-terminus of the second cassette and a 21-amino-acid sequence at the C-terminus of the first cassette. Although these additions are located at the channel extremities, they cannot be assumed to be functionally neutral. Indeed, we observed that the extra proline residue rescued the trafficking defect associated with the T74R variant. Although this finding cannot be generalized to other variants, it raises concerns about the reliability of previously reported trafficking phenotypes for hERG variants characterized using tagged constructs. Interestingly, these proline insertions also induced changes in the biophysical properties of the WT (half-inactivation potential) and T74R (deactivation kinetics), a variant that did not originally affect this property in heterozygous condition. Thus, the introduction of additional sequences at the channel termini represents a second potential source of bias in the functional assessment of *KCNH2* variants and, consequently, in the interpretation of their pathogenicity.

Fourth, our findings identify the co-electroporation of two independent, otherwise identical plasmids - one encoding WT hERG and the other encoding the variant, with neither construct carrying an additional tag - as the most appropriate approach among those investigated. This strategy allows multiple plasmids to enter individual cells, potentially enabling the WT-to-variant expression ratio to be adjusted through the relative amounts of plasmid introduced. It therefore offers a means of more closely reproducing the physiological balance of allelic expression while preserving the native amino acid sequences of both channels. These advantages make co-electroporation a particularly attractive strategy for the functional characterization of *KCNH2* variants, especially when dominant-negative effects are being investigated.

Beyond the characterization of *KCNH2* variants, our findings may also be relevant to other potassium channels that assemble as heterotetramers. For example, *KCNQ1* has been reported to exhibit balanced biallelic expression in the postnatal human heart ^32^, suggesting that similar considerations apply to the functional assessment of its variants. More broadly, an RNA-sequencing analysis of allele-specific expression in primary CD4+ T lymphocytes reported balanced allelic expression for approximately 95.4% of the genes examined ^33^. Although allelic expression patterns may vary across tissues and biological contexts, these findings underscore the potential importance of reproducing physiological allelic expression ratios when developing experimental systems for variant characterization. Ensuring an appropriate balance between WT and variant expression may therefore be critical not only for the interpretation of *KCNH2* variants but also for the reliable functional assessment of genetic variants across a broader range of disease-associated ion channels.

