## Supplemental figures for "Expression-System Biases Compromise the Functional Assessment of *KCNH2* Variants and Clinical Risk Assessment"

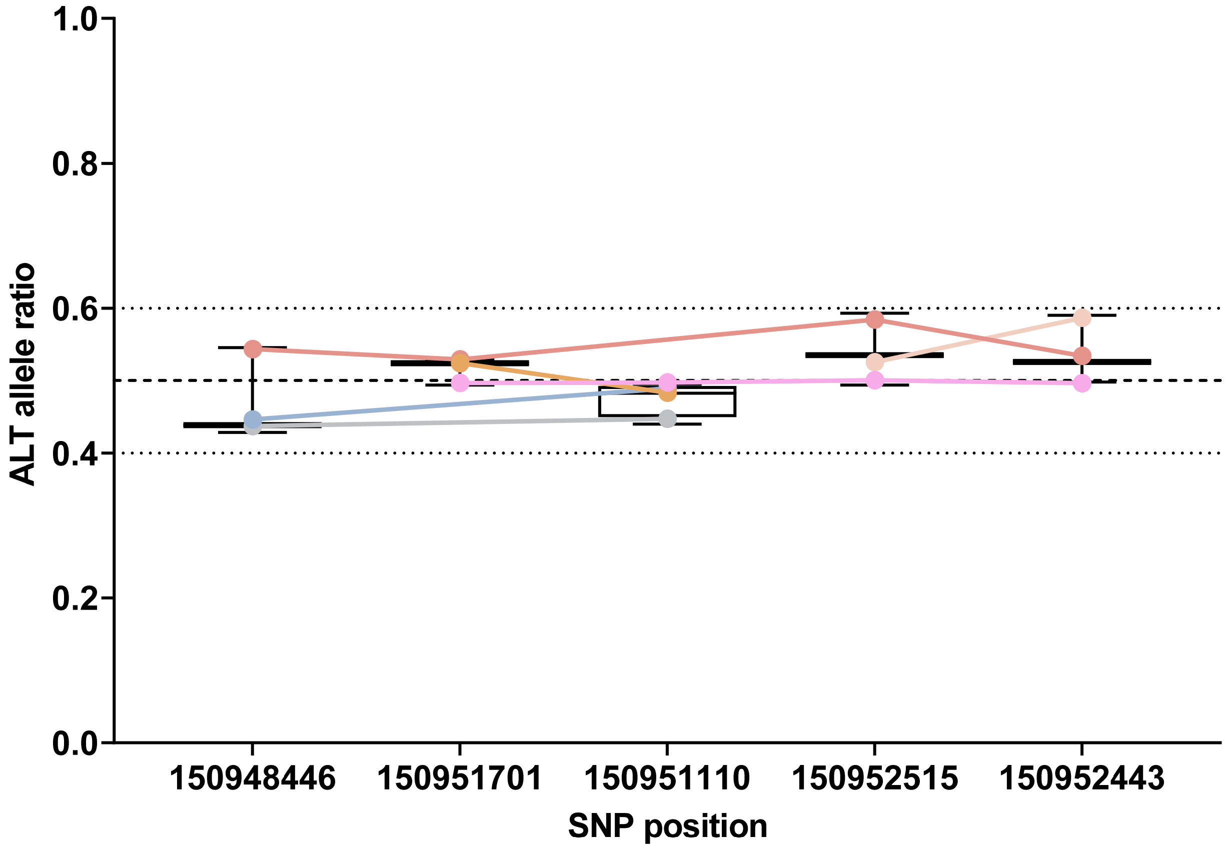


**Supplemental Figure 1**: **Allele-specific expression of *KCNH2* in the GSE298265 RNA-seq dataset.** Alternative allele fractions (ALT/[REF + ALT]) are shown for individual SNPs across samples. Only SNP-sample observations with at least 25 total allele counts were included. Colored points represent individual samples, with lines connecting SNP measurements from the same sample. Boxplots show the distribution of allelic fractions at each SNP. The dashed horizontal line indicates balanced allelic expression (0.5), while dotted lines indicate reference values of 0.4 and 0.6 [1]

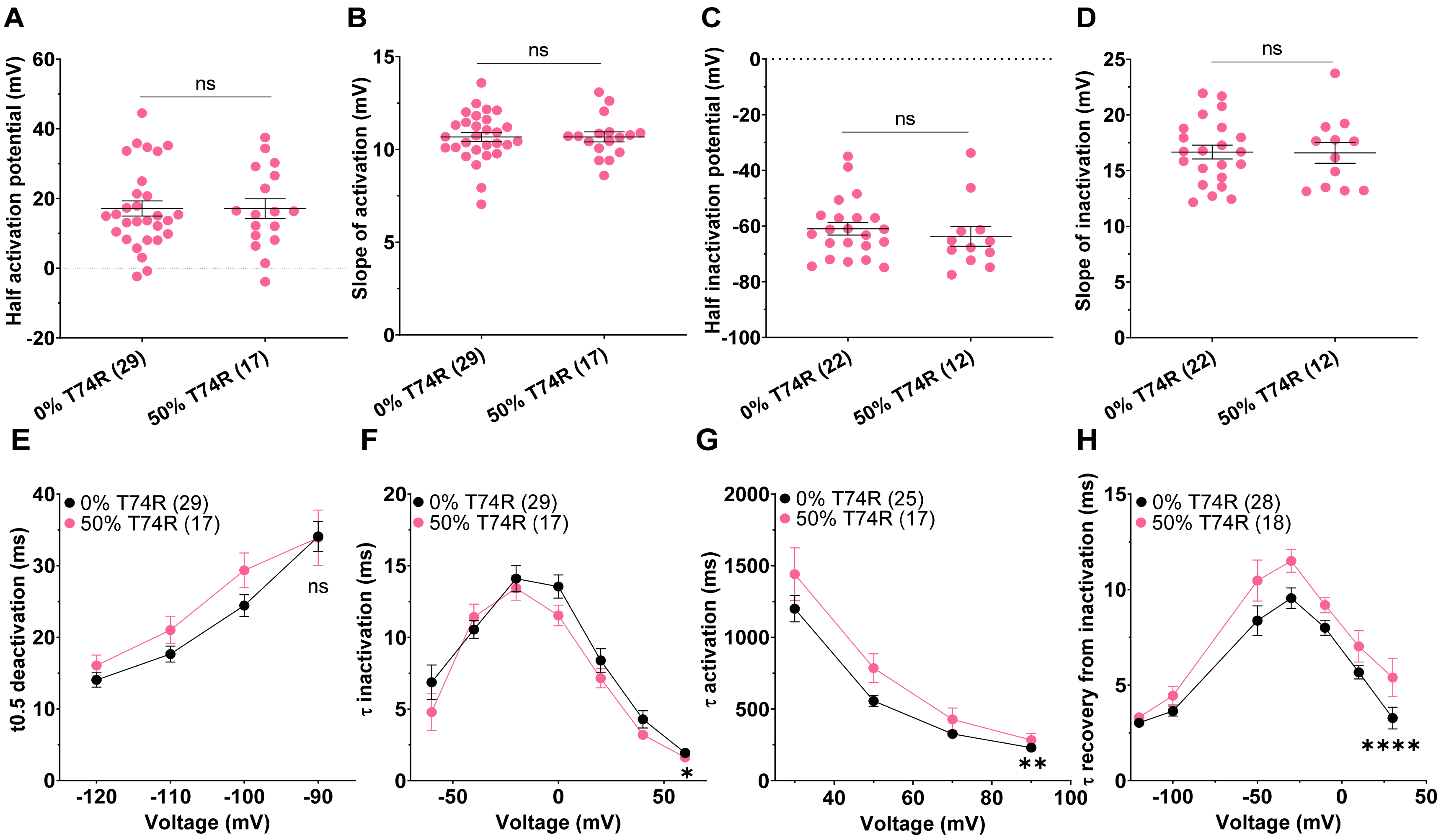


**Supplemental Figure 2**: **Biophysical properties of hERG currents analyzed in 0% and 50% T74R condition using co-transfected plasmids.** A-B) Half activation potential and activation slope. Data are presented as mean ± SEM. C-D) Half inactivation potential and inactivation slope. Data are presented as mean ± SEM. E) Mean ± SEM of half deactivation time for the indicated membrane potentials. F) Mean ± SEM of inactivation time constant for the indicated membrane potentials. ns, not significant; *p value *versus* 0% T74R <.05; two-way ANOVA test. G) Mean ± SEM of activation time constant for the indicated membrane potentials. **p value *versus* 0% T74R <.01; two-way ANOVA test H) Mean ± SEM of recovery from inactivation time constant for the indicated membrane potentials. ****p value *versus* 0% T74R <.0001; two-way ANOVA test.

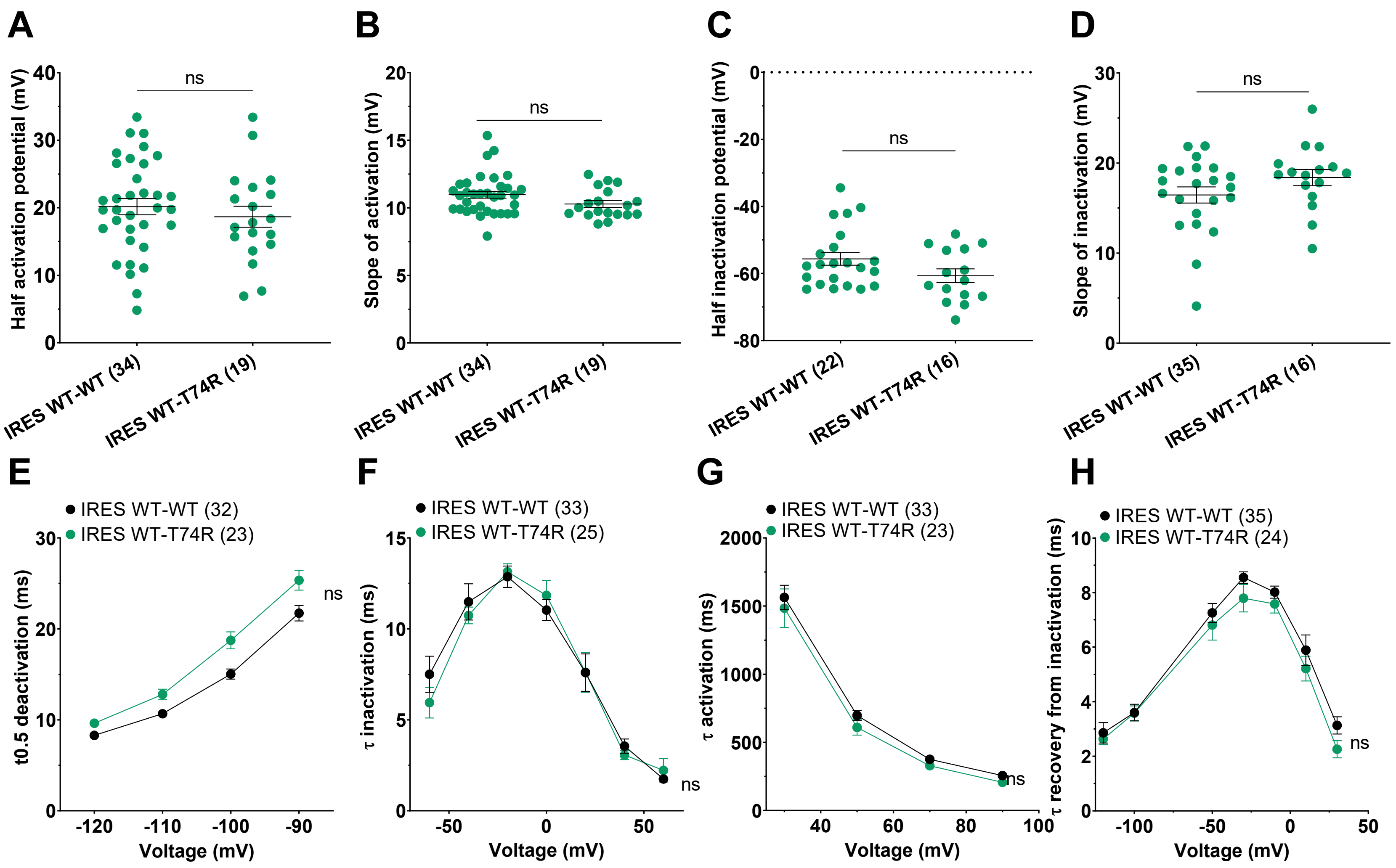


**Supplemental Figure 3**: **Biophysical properties of hERG currents analyzed in condition with detectable currents using IRES plasmids.** A-B) Half activation potential and activation slope. Data are presented as mean ± SEM. C-D) Half inactivation potential and inactivation slope. Data are presented as mean ± SEM. E) Mean ± SEM of half deactivation time for the indicated membrane potentials. F) Mean ± SEM of inactivation time constant for the indicated membrane potentials. G) Mean ± SEM of activation time constant for the indicated membrane potentials. H) Mean ± SEM of recovery from inactivation time constant for the indicated membrane potentials. ns, not significant;

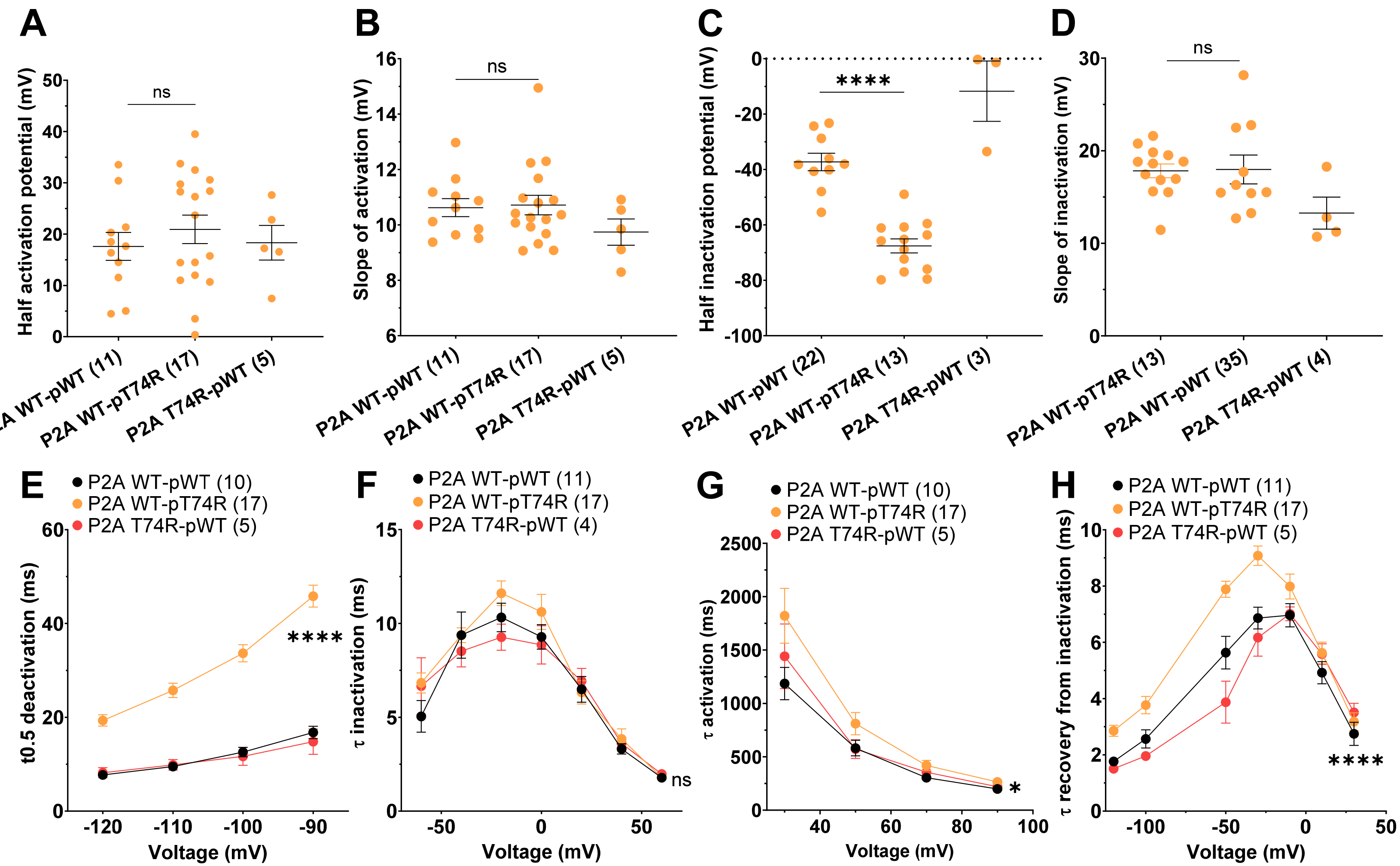


**Supplemental Figure 4**: **Biophysical properties of hERG currents analyzed in condition with detectable currents using P2A plasmids.** A-B) Half activation potential and activation slope. Data are presented as mean ± SEM. ****p<0.0001; Mann-Whitney t-tests C-D) Half inactivation potential and inactivation slope. Data are presented as mean ± SEM. E) Mean ± SEM of half deactivation time for the indicated membrane potentials. ****p value *versus* WT-pWT hERG <.0001; two-way ANOVA test F) Mean ± SEM of inactivation time constant for the indicated membrane potentials. G) Mean ± SEM of activation time constant for the indicated membrane potentials. ns, not significant; *p value *versus* WT-pWT hERG <.01; two-way ANOVA test. H) Mean ± SEM of recovery from inactivation time constant for the indicated membrane potentials. ****p value *versus* WT-pWT hERG <.0001; two-way ANOVA test.

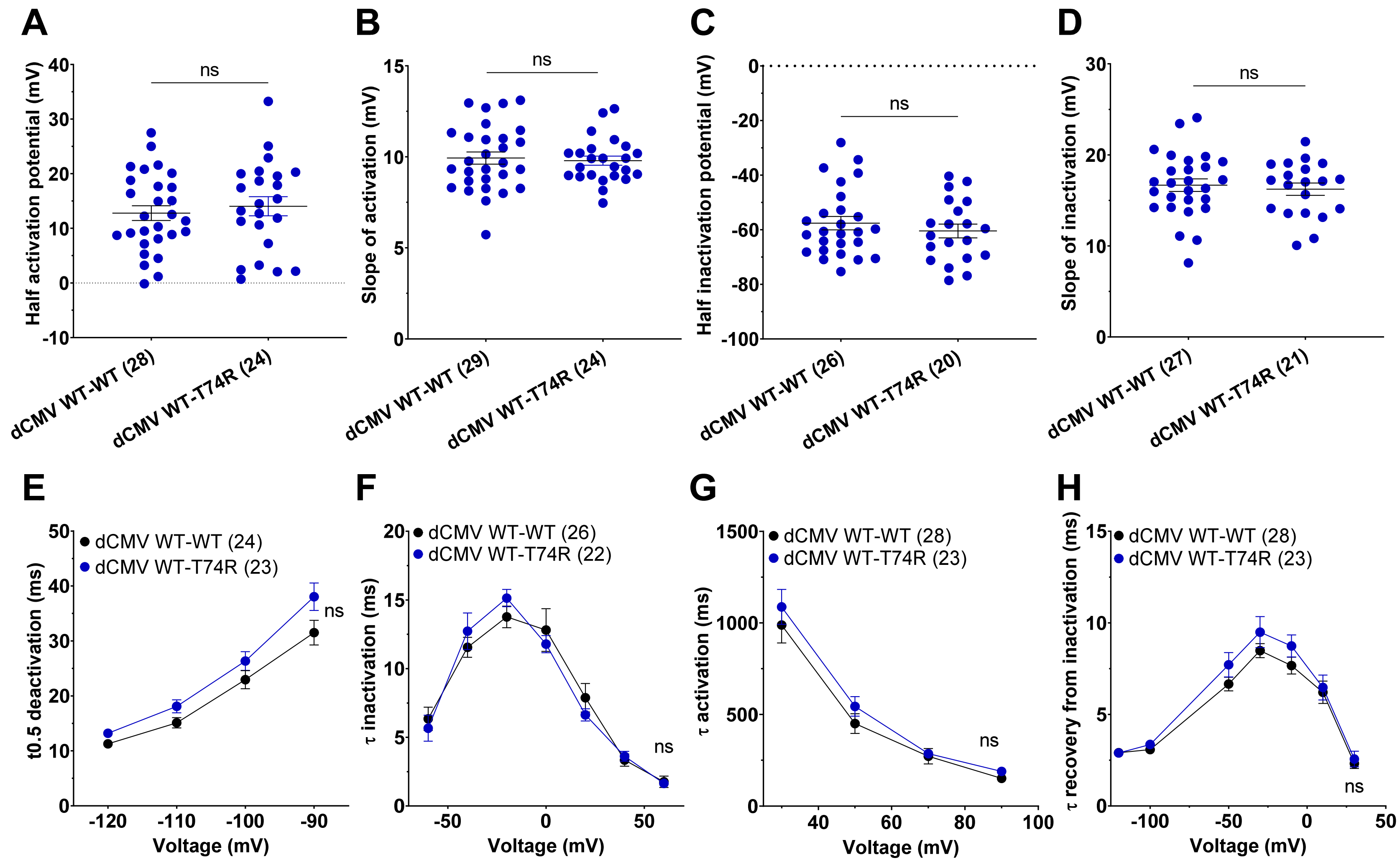


**Supplemental Figure 5**: **Biophysical properties of hERG currents analyzed in condition with detectable currents using dual CMV plasmids transiently expressed.** A-B) Half activation potential and activation slope. Data are presented as mean ± SEM. C-D) Half inactivation potential and inactivation slope. Data are presented as mean ± SEM. E) Mean ± SEM of half deactivation time for the indicated membrane potentials. F) Mean ± SEM of inactivation time constant for the indicated membrane potentials. G) Mean ± SEM of activation time constant for the indicated membrane potentials. H) Mean ± SEM of recovery from inactivation time constant for the indicated membrane potentials. ns, not significant;

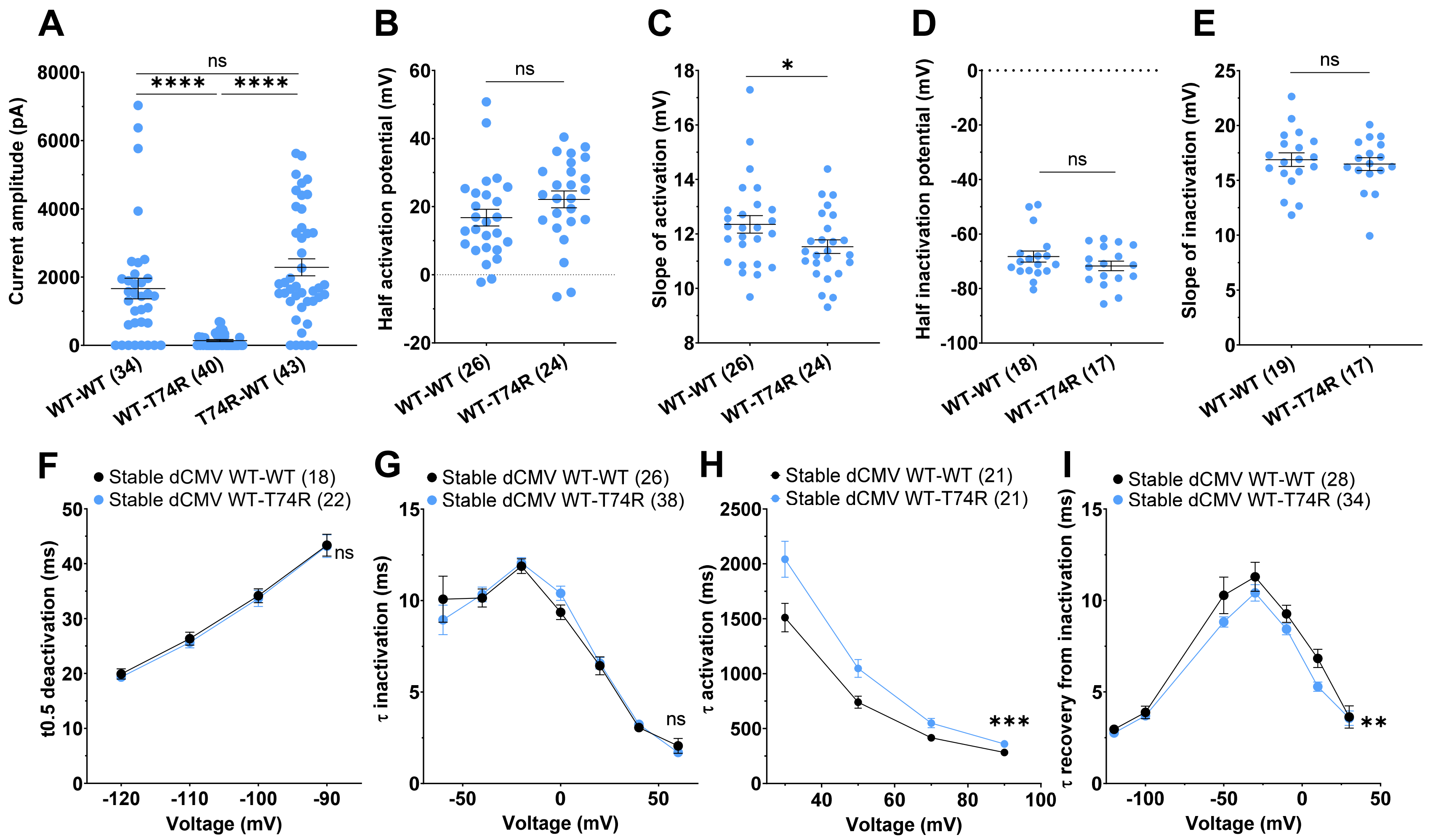


**Supplemental Figure 6**: **Biophysical properties of hERG currents analyzed in condition with detectable currents using dual CMV plasmids stably expressed.** A) Current amplitude of different dCMV stable cell line Data are presented as mean ± SEM . ****p<0.0001; Mann-Whitney t-tests. B-C) Half activation potential and activation slope. Data are presented as mean ± SEM. ns, not significant; *p<0.05; Mann-Whitney t-tests D-E) Half inactivation potential and inactivation slope. Data are presented as mean ± SEM. F) Mean ± SEM of half deactivation time for the indicated membrane potentials. G) Mean ± SEM of inactivation time constant for the indicated membrane potentials. H) Mean ± SEM of activation time constant for the indicated membrane potentials. ***p value *versus* stable dCMV WT-WT hERG <.001; two-way ANOVA test H) Mean ± SEM of recovery from inactivation time constant for the indicated membrane potentials. **p value *versus* WT-WT hERG <.01; two-way ANOVA test.
